# Circulating miR-30b-5p is upregulated in Cavalier King Charles Spaniels affected by early myxomatous mitral valve disease

**DOI:** 10.1101/2022.03.17.484775

**Authors:** Mara Bagardi, Sara Ghilardi, Valentina Zamarian, Fabrizio Ceciliani, Paola G. Brambilla, Cristina Lecchi

## Abstract

There is a growing interest in developing new molecular markers of heart disease in young Cavalier King Charles Spaniels affected by myxomatous mitral valve disease. The aim of the study was to measure the abundance of 3 circulating microRNAs and their application as potential biomarkers in the plasma of Cavalier King Charles Spaniels with early asymptomatic myxomatous mitral valve disease. 33 dogs affected by the disease in American College of Veterinary Internal Medicine (ACVIM) stage B1 were divided in three groups (11 younger than 3 years, 11 older than 3 years and younger than 7 years, and 11 older than 7 years), and 11 healthy (ACVIM stage A) Cavalier King Charles Spaniels were included as the control group. This is a prospective cross-sectional study. The abundance of three circulating microRNAs (miR-1-3p, miR30b-5p, and miR-128-3p) was measured by quantitative real-time PCR using TaqMan® probes. Diagnostic performance was evaluated by calculating the area under the receiver operating curve (AUC). miR-30b-5p was significantly higher in ACVIM B1 dogs compared to ACVIM A subjects, and the area under the receiver operating curve was 0.79. According to the age of dogs, the abundance of miR-30b-5p was statistically significantly higher in group B1<3y (2.3 folds, *P* = 0.034), B1 3-7y (2.2 folds, *P* = 0.028), and B1>7y (2.7 folds, *P* = 0.018) than in group A. The area under the receiver operating curves were fair in discriminating between group B1<3y and group A (AUC 0.780), between B1 3-7y and A (AUC 0.78), and good in discriminating between group B1>7y and A (AUC 0.822). miR-30b-5p changed in the plasma of dogs at the asymptomatic stage of disease, particularly at a young age.

## Introduction

Myxomatous mitral valve disease (MMVD) is a cardiovascular disease affecting dogs, which progress from mitral regurgitation (MR) to eventually heart failure. The disease causes about 10% of all the deaths in this species [1]. Although MMVD seems to be a genetic disorder, a mutation has not yet been identified [2]. The incidence is age-related and is particularly high in some breeds such as the Cavalier King Charles Spaniel (CKCS). In fact, half of the CKCSs are estimated to be affected by MMVD at the age of 6-7, while at 10 years of age almost all of them are [1,3-5]. Evidence from highly susceptible breeds such as CKCS and Dachshund shows a strong inherited component to the disease and suggests a polygenic inheritance [2,6,7]. Due to the lack of early signs, symptoms, and predictive biomarkers, early diagnosis is difficult. Identifying reliable specific biomarkers is desirable, especially for screening and breeding programs. In human medicine, microRNAs (miRNAs) are potentially suitable markers of cardiovascular diseases [8,9]. miRNAs exert their function by repressing the translation of target genes, and by regulating protein production through different mechanisms in several pathophysiological conditions, including myocardial infarction, hypertrophy, fibrosis, and inflammation. MiRNAs can be secreted into extracellular fluids, including plasma and serum, within vesicles, such as exosomes, or in conjunction with lipoproteins and RNA-binding proteins, namely Argonaute (AGO). By doing so, they are relatively stable even under tough conditions such as long-time storage at room temperature, and low or high pH [10-14]. Aberrant expression of miRNAs is associated with several human and veterinary disorders, including cancer and heart diseases [15-22]. The dysregulation of circulating miRNAs was previously investigated also in MMVD-affected dogs following different approaches, including quantitative real-time PCR (RT-qPCR), microarray, and next-generation sequencing (NGS) [23-28]. Most of the dogs enrolled in these studies were classified following American College of Veterinary Internal Medicine (ACVIM) guidelines as stage C and D, while only one study performed analysis also on dogs older than 8 years in ACVIM stages B1 and B2 [26-29].

The present study aimed at improving MMVD assessment in CKCSs at the asymptomatic ACVIM stage B1 by ascertaining how three miRNAs, previously associated with MMVD, are modulated in the plasma of CKCSs divided according to their age at the time of diagnosis (younger than 3 years, between 3 and 7 years, and older than 7 years). Thereby, miRNAs are investigated for their potential use as biomarkers to identify asymptomatic dogs in ACVIM stage B1. The decision to focus the study on this ACVIM class was driven by the fact that these dogs are most subjected to breed screening, and therefore are targeted as potential breeders. An early diagnosis of MMVD is only achievable through echocardiographic examination. However, the disease goes undetected in subjects that have no clinical signs and that do not present heart murmurs at the clinic visit. Currently, there are no tests available to outline the evolution of this disease in the CKCS, therefore the aim of this study is to cover this gap by investigating miRNAs as potential biomarkes for early diagnosis of MMVD. Highlight the risk of the development of the disease at an earlier stage will favour a preventive screening and a mitigating therapeutical approach.

## Material and methods

### Clinical and echocardiographic examinations

The study included 44 owned CKCSs visited at the Cardiology Unit of the Department of Veterinary Medicine, University of Milan, between May 2019 and July 2020. Informed consent was signed by the owners, according to the ethical committee statement of the University of Milan, number 2/2016, and a high standard of care was provided throughout each examination.

During a routine veterinary visit, a cardiological evaluation was performed in dogs fasted for at least 12 hours. The clinical data of the animals included animal history, and clinical and echocardiographic examinations. The cardiovascular system was evaluated by checking the presence/absence of murmurs by two different well-trained operators, respectively a third year PhD student in cardiology and a professor with more than twenty years of practice in clinical veterinary cardiology (MB and PGB). The evaluated auscultatory findings included presence/absence, timing, and intensity of the murmur (0 = absent; 1 = I-II/VI left apical systolic or soft; 2 = III-IV/VI bilateral systolic or moderate and loud respectively; 3 = V-VI/VI bilateral systolic or palpable) [30]. Blood pressure was indirectly measured with a Doppler method according to the ACVIM consensus statement [31,32]. Peripheral venous blood sampling was performed at the end of the examination. Blood was collected from the jugular or cephalic vein in two 2.5-mL EDTA tubes.

The echocardiographic exam was used to diagnose MMVD. A standard transthoracic echocardiographic examination was performed with My Lab50 Gold Cardiovascular ultrasound machine (Esaote, Genova, Italy), equipped with multi-frequency phased array probes (3.5-5 and 7.5-10 MHz), chosen according to the weight of the subject. Videoclips were acquired and stored using the echo machine software for offline measurements. The exam was performed by a certified cardiologist (MB) according to a standard procedure with concurrent continuous electrocardiographic monitoring [33]. All examinations were performed without pharmacological restraint. Dogs were classified according to the ACVIM classification scheme [29].

Inclusion criteria for dogs in the clinically normal group (ACVIM A) were: no echocardiographic evidence of heart disease, no clinical signs, no abnormalities on results of a complete blood count and biochemical analyses, and no history of medical treatment within the previous 6 months. Inclusion criteria for dogs with MMVD at stage B1 were: echocardiographic evidence of a thickened or prolapsed mitral valve and mitral valve regurgitation, no evidence of left atrial dilatation, defined as a left atrial-to-aortic root ratio (LA/Ao) <1.6 on 2-dimensional echocardiography, and no left ventricle dilation, defined as left ventricular normalized dimensions in diastole (LVIDdN) <1.7. Dogs that presented left atrial and/or ventricular remodeling, but not severe enough to meet the current guidelines criteria for ACVIM class B2, were also included [29]. The degree of MR (jet size) was assessed using color Doppler and calculating the maximal ratio of the regurgitant jet area signal to the left atrium area (ARJ/LAA ratio) [34]. The regurgitant jet size was estimated as the percentage of the left atrial area (to the nearest 5%) that was occupied by the larger jet, and it was considered as trivial or trace (<10%), mild (between 10 and 30%), moderate (between 30 and 70%) or severe (>70%) [34,35]. Mitral regurgitation was considered as trivial when the regurgitant jet was not detectable in all systolic events, while it was considered as trace when it was always visible [35]. Four groups of 11 client-owned dogs were included in the present study: group A, or heathy control, group B1<3 with dogs younger than 3 years; group B1 3-7, with dogs older than 3 years and younger than 7 years, and B1>7 with dogs older than 7 years [36,37].

Dogs with asymptomatic MMVD and cardiac remodeling (ACVIM stages B2), dogs with symptomatic MMVD (ACVIM stages C and D), or with other systemic diseases such as systemic hypertension, uncontrolled hypothyroidism, hyperadrenocorticism, primary pulmonary hypertension, neoplasia, and other cardiac abnormalities such as dilated cardiomyopathy, congenital cardiac abnormalities, endocarditis, and severe arrhythmia were excluded from the study.

### Small RNA isolation and RT-qPCR quantification

Blood samples for small RNA isolation were collected in 2.5 ml EDTA-K3 tubes. Within 2 hours, the samples were centrifuged at 800 g for 15 minutes. Plasma was stored at – 80°C until RNA isolation.

Small RNA was extracted using the miRNeasy Serum/Plasma Kit (Qiagen, catalogue number 217184, Milan, Italy). An aliquot of 150 µL of plasma per sample was thawed on ice, and centrifuged at 3000 × g for 5 min at 4°C. RNA was extracted using miRNeasy Serum/Plasma Kits (Qiagen, Cat. No. 217184, Milano, Italy) following the manufacturer’s instructions. One mL of Qiazol (Qiagen) was added to an aliquot of 150 μl of plasma per sample. After incubation at room temperature for 5 min, 25 fmol of the exogenous synthetic spike-in control *Caenorhabditis elegans* miRNA cel-miR-39 (Qiagen, Cat. No. 219610) was spiked into samples at the beginning of the extraction procedure to check both the extraction of miRNAs and the efficiency of the cDNA synthesis. RNA extraction was then carried out according to the manufacturer’s instructions. RNA yield, as well as successful RNA purification without contaminations of proteins or residues from the isolation procedure, is was assayed using 1 μl of eluted RNA applied to a NanoDrop ND-1000 spectrophotometer. The 260/280 nm ratio was between 1.8 and 2.2 for all RNA samples, and the range of 260/230 nm ratio was from 2.0 to according to MIQE guidelines [38,39]. To obtain cDNA, reverse transcription was performed on 10 ng of total RNA using a TaqMan Advanced miRNA cDNA Synthesis Kit (Cat. No. A28007, Applied Biosystems) following the manufacturer’s instructions.

RT-qPCR was performed following the MIQE guidelines [38,39]. The small RNA TaqMan assays were performed according to the manufacturer’s instructions using the selected primer/probe assays (ThermoFisher Scientific), which are also specific for canine miRNAs, including: cel-miR-39-3p (assay ID 478293_mir); miR-1-3p (assay ID 477820_mir) [28]; miR-30b-5p (assay ID 478007_mir) [40]; miR-128-3p (assay ID mmu480912_mir) [28]. The reference miRNA was miR-16-5p (assay ID rno481312_mir). Quantitation was performed on 15 µl in a CFX Connect Real-Time PCR Detection System (Bio-Rad) using 7.5 μl of 2X TaqMan Fast Advanced Master Mix (Cat. No. 4444557), 0.75 μl of miRNA specific TaqMan Advanced assay reagent (20X), 1 μl of cDNA, and water to make up the remaining volume. The thermal cycling profile was as follows: 50°C for 2 min, 95°C for 3 min, 40 cycles at 95°C for 15 s and 60°C for 40 s. No-RT controls and no-template controls were included. MicroRNA expressions are presented in terms of fold change normalized to miR-16 as reference miRNA, and sample A as reference sample using the formula 2^−ΔΔCq^ on Bio-Rad CFX Maestro Software [39].

### Statistical analysis

Statistical analysis was performed using XLStat software for Windows (Addinsoft, New York, USA). Data were tested for normality using the Shapiro–Wilk test; when the data were not normally distributed, the nonparametric Kruskal-Wallis test was applied. Receiver operating characteristic (ROC) analysis was performed and the area under the ROC curve was considered as a measure of the diagnostic accuracy using the definition suggested by Šimundić in 2009 [41]. The diagnostic value was calculated for miRNA that showed significant differential expression in the canine blood. Statistical significance was accepted at a *P* value ≤ 0.05 and all the significance values were adjusted according to the Bonferroni post-hoc correction.

## Results

### Demographics and characteristics of study subjects

The median age of the 44 included CKCSs was 3.3 years (IQR_25-75_ 1.81-6.99), and the median body weight was 8.1 Kg (IQR_25-75_ 7.48-9.68). 14 subjects (31.82%) were males, and 30 (68.18%) were females. Study population characteristics (clinical and echocardiographic data), grouped according to the ACVIM classes, and, for the B1 class, to the age at the time of MMVD diagnosis, are shown in Table 1. Weight was lower in B1<3 (*P* = 0.040) and A (*P* = 0.029) subjects compared with the B1>7 group, whereas echocardiographic variables were not statistically different among age groups (*P* > 0.05).

**Table 1.** Clinical and echocardiographic data of all included CKCSs divided according to ACVIM classification scheme and age at the time of the MMVD diagnosis for subjects belonging to ACVIM stage B1.

|  | **Overall population** | **A** | **B1<3** | **B1 3-7** | **B1>7** |
| --- | --- | --- | --- | --- | --- |
| **Number of dogs** | 44 | 11 | 11 | 11 | 11 |
| **Sex** | 30F (8NF)<br>14M | 8F (1NF)<br>3M | 6F<br>5M | 6F (1 NF)<br>5M | 10F (6 NF)<br>1M |
| **Age (years)** | 3.30 (1.81-6.99) | 1.96 (1.73-2.88) | 1.52 (1.07-2.21) | 3.88 (3.49-4.30) | 8.14 (7.66-8.68) |
| **Weight (kg)** | 8.10 (7.48-9.68) | 7.75 (7.25-7.95)<sup>a</sup> | 7.80 (6.83-8.00)<sup>a</sup> | 9.40 (7.63-9.88) | 10.00 (9.35-10.40) |
| **SBP (mmHg)** | 135 (110-145) | 125 (115-135) | 130 (120-140) | 130 (120-140) | 140 (125-150) |
| **Murmur** | 29 grade 0<br>10 grade 1<br>5 grade 2 | 11 grade 0 | 10 grade 0<br>1 grade 1 | 8 grade 0<br>3 grade 1 | 6 grade 1<br>5 grade 2 |
| **Regurgitant jet size** | 11 grade 0<br>5 grade 1<br>8 grade 2 | 11 grade 0 | 5 grade 1<br>6 grade 2 | 2 grade 2<br>9 grade 3 | 6 grade 3<br>5 grade 4 |
|  | 15 grade 3 |  |  |  |  |
|  | 5 grade 4 |  |  |  |  |
| **ESVI ml/m<sup>2</sup>** | 17.95 (14.55-25.93) | 16.60 (11.19-17.95) | 16.69 (14.73-21.67) | 19.48 (15.73-26.03) | 22.15 (17.29-27.93) |
| **EDVI ml/m<sup>2</sup>** | 56.15 (48.28-71.91) | 50.50 (45.54-61.87) | 51.99 (41.97-62.64) | 56.38 (52.42-85.29) | 68.40 (58.98-76.59) |
| **LA/Ao** | 1.15 (1.08-1.26) | 1.21 (1.09-1.36) | 1.08 (1.02-1.20) | 1.17 (1.12-1.19) | 1.15 (1.08-1.29) |
| **E (m/s)** | 0.73 (0.66-0.80) | 0.76 (0.71-0.84) | 0.68 (0.61-0.79) | 0.70 (0.67-0.75) | 0.68 (0.67-0.81) |
| **E/A** | 1.30 (1.18-1.43) | 1.29 (1.21-1.37) | 1.47 (1.36-1.60) | 1.21 (1.16-1.45) | 1.22 (0.99-1.34) |
| **EF (%)** | 67 (58-73) | 67 (62-77) | 62 (59-70) | 68 (57-74) | 67 (57-71) |
| **FS (%)** | 35 (30-40) | 35 (32-43) | 31 (30-38) | 36 (29-41) | 37 (28-39) |
| **LVIDas** | 0.84 (0.77-0.95) | 0.82 (0.70-0.84) | 0.82 (0.78-0.90) | 0.87 (0.79-0.96) | 0.90 (0.81-0.98) |
| **LVIDad** | 1.36 (1.29-1.51) | 1.31 (1.26-1.42) | 1.33 (1.22-1.43) | 1.37 (1.33-1.62) | 1.48 (1.40-1.55) |

E = E wave velocity; E/A = E and A waves ratio; EDVI = end diastolic volume index; EF = ejection fraction; ESVI = end systolic volume index; FS = shortening fraction; LA/Ao = left atrium to aortic root ratio; LVIDad = left ventricular normalized dimensions in diastole; LVIDas = left ventricular normalized dimensions in systole; Murmur = left systolic heart murmur intensity: 0 = absent, 1 = I-II/VI left apical systolic, or soft, 2
= III-IV/VI bilateral systolic, or moderate and loud; Sex: F = female, NF = neutered female, M = male; SBP = systemic blood pressure; Regurgitant jet size: 0 = absent, 1 = trivial, 2 = trace, 3 = mild, 4 = moderate. All data are expressed as median and IQR25-75 ranges. aValues within a row differ significantly at $P < 0.05$ from B1>7 subjects.

### miR-30b-5p is differentially expressed in MMVD-affected dogs

Small RNA was extracted from plasma, and the spike-in cel-miR-39 was quantified in all collected samples exhibiting a mean Cq of 26.09 (SD 1.13). Three miRNAs, namely miR-1-3p, miR-30b-5p, and miR-128-3p, were detected in all plasma samples (Figs 1A-F). The comparative analysis demonstrated that one miRNA, namely miR-30b-5p, had a significant differential abundance in the plasma of MMVD-affected dogs compared to the healthy group. In detail, the abundance of miR-30b-5p increased 2.4 folds (*P* = 0.006) in group B1 compared to group A (Fig 1B). When group B1 was further split according to the age of dogs, the expression of miR-30b-5p remained significantly higher (Fig 1E): group B1<3 (2.3 folds *P* = 0.034), B1 3-7 (2.2 folds *P* = 0.028), and B1>7 (2.7 folds *P* = 0.018) showed a higher level of miR-30b-5p than group A. No differences were found in the amount of miR-1-3p (Figs 1A and D) and miR-128-3p (Figs 1C and F). The age proved not to be correlated with the expression of the analyzed miRNAs, neither in the entire population nor in each age class (*P* > 0.05).

**Fig 1.**
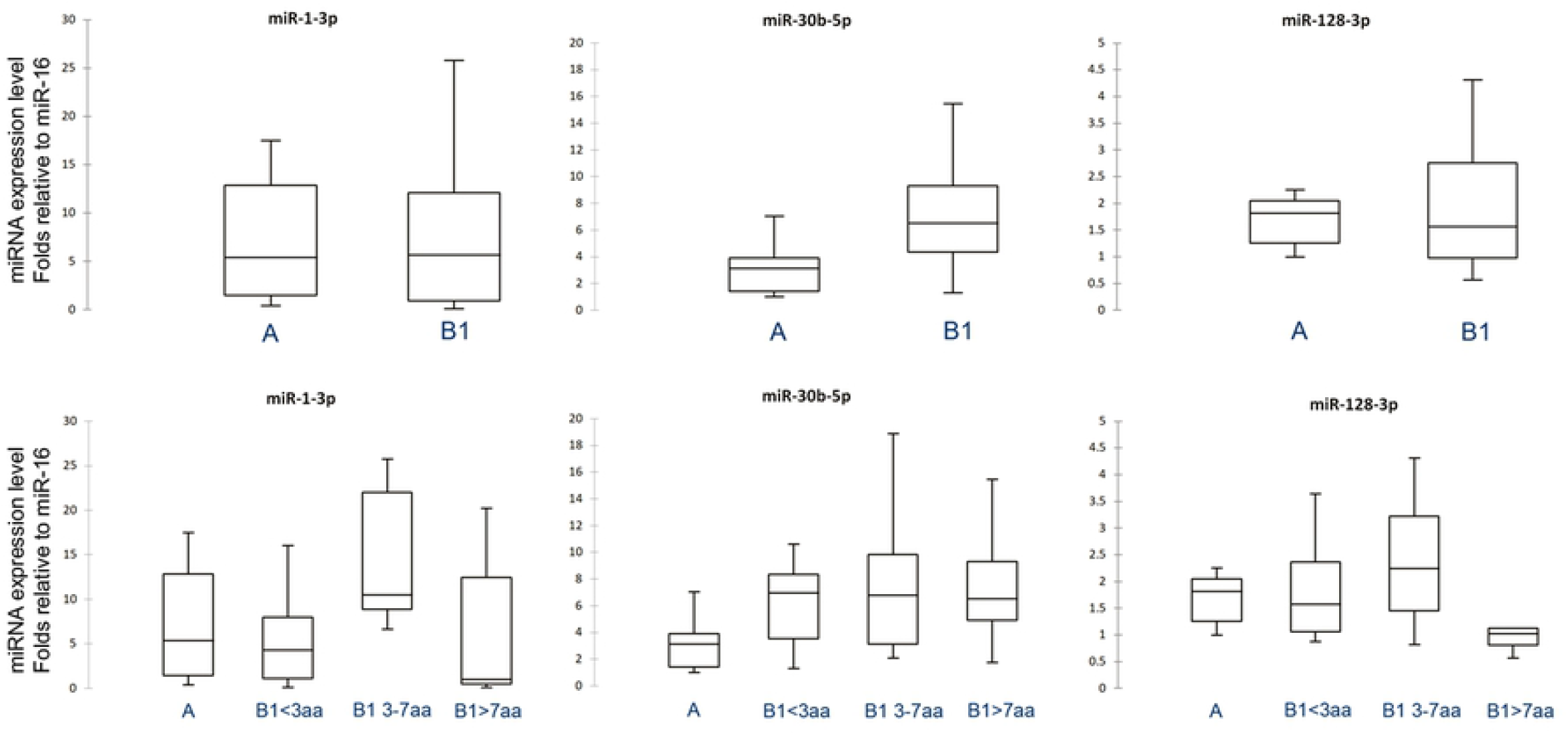
Expression levels of miR-1-3p, miR-30b-5p, and miR-128-3p between groups. Expression levels of the three miRNAs between groups A and B1 (Figs 1A-C, respectively), and between A and B1 divided according to age at MMVD diagnosis (Figs 1D-F, respectively). miR-30b-5p increased 2.4 folds (*P* < 0.05) in group B1 compared to A (Fig 1B). Splitting group B1 according to the age of dogs, the expression of miR-30b-5p remained significantly higher (Fig 1E). Group B1<3 (2.3 folds, *P* = 0.034), B1 3-7 (2.2 folds, *P* = 0.028), and B1>7 (2.7 folds, *P* = 0.018) expressed a higher level of miR-30b-5p than group A. No differences were found in the amount of miR-1-3p (Figs 1A and 1D) and miR-128-3p (Figs 1C and 1F).

### Diagnostic performance of miR-30b-5p discriminating between MMVD-affected and healthy dogs

To evaluate the diagnostic value of miR-30b-5p in plasma, ROC curve analysis was performed, and the associated AUC was used to confirm the diagnostic potency. Cut-off points were set to maximize the sum of sensitivity and specificity. The ability of miR-30b-5p to separate the tested samples into healthy (stage A) or MMVD-affected (stage B1) was defined as “diagnostic accuracy” and was measured by the area under the curve (AUC). miR-30b-5p proved to be efficient in discriminating between groups A and B1 (AUC = 0.79; 95% CI 0.65-0.93) (Fig 2A). Even after dividing group B1 according to age, it could efficiently discriminate between group A and group B1<3 (AUC = 0.78; 95% CI 0.60-0.96) and group A and group B1 3-7 (AUC = 0.78; 95% CI 0.60-0.96) (Figs 2B and 2C, respectively), but in particular it proved to be really effective in discriminating group A from B1>7 (AUC = 0.82; 95% CI 0.65-0.99) (Fig 2D) (Table 2). Thus, miR-30b-5p can discriminate between healthy (stage A) and asymptomatic MMVD-affected dogs (stage B1).

**Table 2.** Area under the curve (95% confidence interval), cut off values, and sensitivity and specificity of miR-30b-5p in CKCSs’ plasma.

|  | AUC | 95% CI | P value | Cut off | Se-Sp |
| --- | --- | --- | --- | --- | --- |
| A vs B1 | 0.793 | 0.653-0.933 | > 0.0001 | 3.98 | 0.759-0.800 |
| **A vs B1<3** | 0.780 | 0.603-0.957 | 0.0019 | 4.52 | 0.800-0.700 |
| **A vs B1 3-7** | 0.780 | 0.600-0.959 | 0.0023 | 4.37 | 0.800-0.700 |
| **A vs B1>7** | 0.822 | 0.655-0.989 | 0.0002 | 4.37 | 0.800-0.889 |

AUC = area under the ROC curve; CI = confidence interval; Se = sensitivity; Sp = specificity.

**Fig 2.**
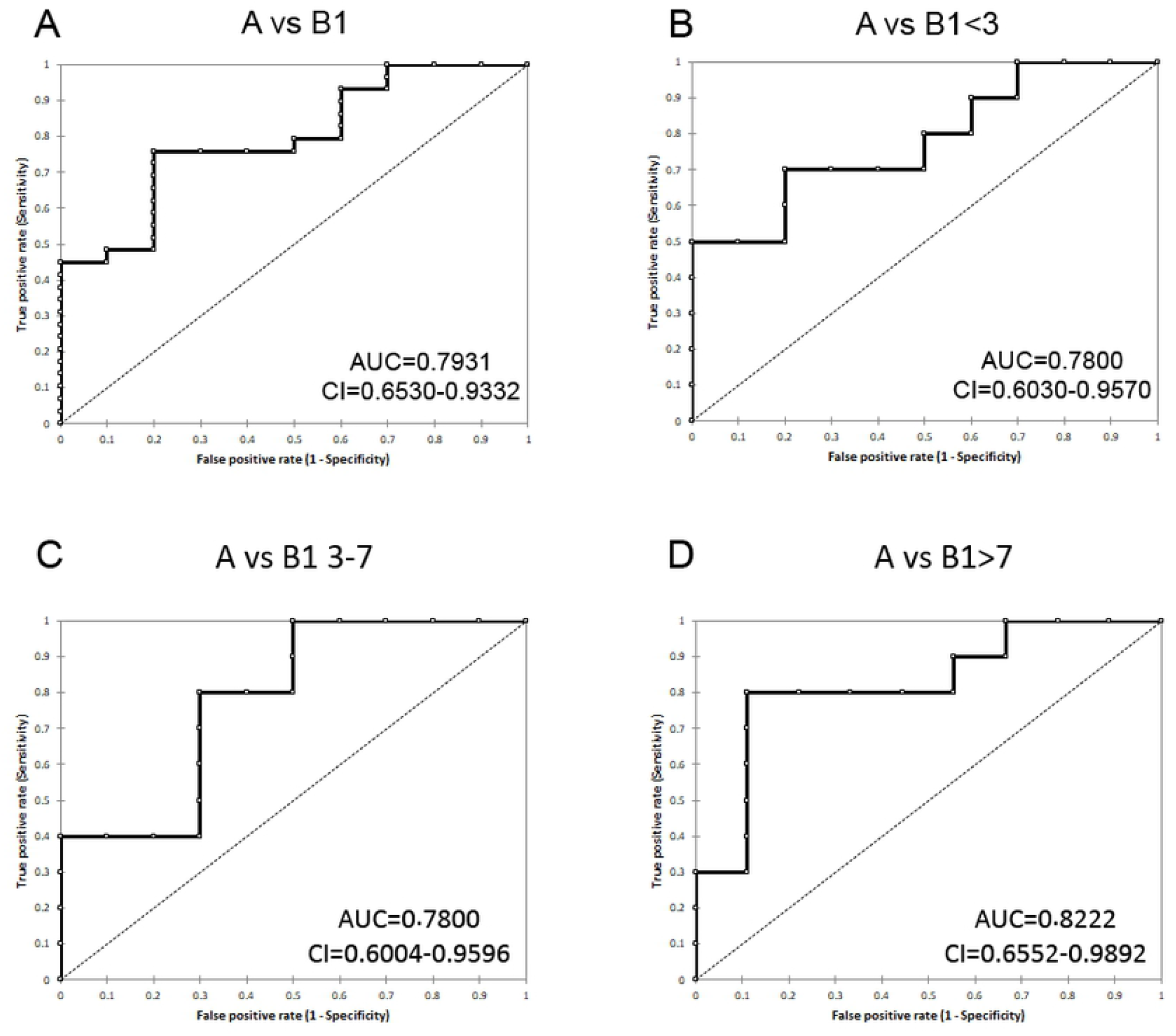
ROC curves for miR-30b-5p. Discrimination capacity between group A and group B1 (Fig 2A), group A and group B1<3 (Fig 2B), group A and group B1 3-7 (Fig 2C), and group A and B1>7 (Fig 2D). miR-30b-5p can discriminate between healthy and asymptomatic MMVD-affected dogs.

## Discussion

The present study reported the relationship between the abundance of circulating miR-30b-5p and the presence of MMVD, a disease often associated with congestive heart failure (CHF), even in young CKCSs. Our results showed that miR-30b-5p is significantly upregulated in asymptomatic MMVD-affected CKCSs (ACVIM stage B1 without cardiac remodeling, or with remodeling changes not severe enough to meet the current guidelines criteria for ACVIM class B2) compared to healthy (ACVIM stage A) dogs. We further demonstrated that miR-30b-5p upregulation is also detectable in young dogs (age <3, ranging from 6 months to 2.4 years), even in MMVD-affected subjects without audible heart murmurs.

Yang and colleagues investigated the cargo of exosomes purified from the plasma of MMVD-affected dogs using an array-based approach, demonstrating that when the False Discovery Rate (FDR) was set at 20%, 78 miRNAs were dysregulated, while with an FDR at 10% no differences were pointed out in either the exosome miRNAs or the whole plasma [25]. Another study, including old dogs (ranging from 8.2 to 13.8 years) with CHF secondary to MMVD (ACVIM stage C), reported that 326 miRNAs were differently modulated comparing healthy (ACVIM stage A) to CHF-affected dogs (ACVIM stage C); the validation step, performed by RT-qPCR, demonstrated the upregulation of miR-133, miR-1, let-7e, and miR-125, and the downregulation of miR-30c, miR-128, miR-142, and miR-423 [28]. Although the study focused on a group of animals affected by a severe disease with clinically detectable signs, the results appeared worthy to be further investigated even in younger patients, prompting us to include miR-1 and miR-128 in the present study. Based on other results previously reported [40] of a study that also included old dogs (range, 10.17 ± 3.36 years), we identified miR-30b-5p as a potential marker to be further investigated in a younger cohort of MMVD-affected CKCSs of ACVIM stage B1.

Since the molecular background of MMVD is not fully elucidated yet, identifying any specific markers (prognostic and/or therapeutic) would be of great clinical value for recognizing asymptomatic patients, especially at a young age. The diagnosis of MMVD is based on the echocardiographic evaluation of the mitral valve and its leaflets’ thickness, which sometimes is hard to quantify since mildly-affected valves work adequately, and the lesions apparently don’t affect hemodynamic, given the absence of cardiac remodeling and clinical signs. Myxomatous mitral valve disease is age-related, and the prevalence in old small-breed dogs is up to 100%, particularly in chondrodystrophic breeds such as Cocker Spaniels, Dachshunds, and Beagles. CKCSs are more susceptible to develop CHF due to MMVD at a younger age than other breeds [42]. Thus, especially in susceptible breeds such as the CKCS, MMVD occurs at a very young age and progresses over time in different and unpredictable grades. MMVD development is regarded as a hereditary character in this breed and has been associated with a multi-factorial polygenic transmission mode: therefore, several genes are involved, and a defined threshold of expression must be reached before the disease occurs [2,5,7].

Although miRNAs are currently intensively investigated in human medicine because of their diagnostic potential in many different conditions, there are only a few reports related to circulating miRNAs studies in dogs affected by MMVD, and no study about the early diagnosis of this disease in a predisposed breed such as the CKCS.

This study identified a biomarker that may have an impact in both implementing preventing programs through genetic selection and in clinical practice. These results confirm that in CKCSs, as already demonstrated in humans, there is a differential expression of miRNAs, suggesting that their expression profiles are distinct for dogs with MMVD compared to healthy dogs. We demonstrated that miR-30b-5p could discriminate among ACVIM stage A CKCSs and ACVIM stage B1 CKCSs younger than 3 years, without heart murmurs, without clinical signs, but with an echocardiographic diagnosis of MMVD. Our results disagree with the previously reported findings on Dachshunds [40], that demonstrated that miR-30b decreased in the plasma of ACVIM stage B subjects compared with ACVIM stage A. This contraddictory result could be explained firstly by the different enrolment strategy (the previous study considered all ACVIM B dogs without distinguishing B1 from B2), and secondly by the different ages of enrolled patients; the present investigation focused mainly on young dogs (33 out of 44 dogs were younger than 4 years), while the previous study only considered old dogs (range, 10.17 ± 3.36 years) [40]. Furthermore, the miR-30 family is abundantly expressed in the heart and its decrease is strictly related to several heart diseases that result in ventricular remodeling [43]. The dogs enrolled in our study did not have ventricular remodeling yet, but only slightly affected valves. Since the miR-30 family exerts antiapoptotic and anti-inflammatory activities [44,45], we hypothesized that the expression of miR-30b may increase during the early stage of MMVD in young CKCSs to protect the cardiomyocytes from inflammation and apoptosis and oppose to the ventricular valves remodeling.

The identification of dogs with early asymptomatic MMVD through the evaluation of miR-30b-5p could help the clinicians and the breeders to better focalized screening programs in this breed and to better select the breeders. Patients with these characteristics should then be subjected to a closer follow-up. For these reasons, miRNAs may be candidates as novel biomarkers and may provide the basis for further investigations, in order to assess the follow-up and characterize the evolution of the disease in the CKCS [46].

This study presents some limitations. The utility of circulating miRNAs as biomarkers of many diseases has attracted considerable attention over recent years. However, it is also worth pointing out that the clinical application of miRNAs as biomarkers is still limited. One of the most significant obstacles is the difficulty concerning the normalization of circulating miRNAs. Spiked in synthetic miRNAs are widely used to normalize serum and plasma miRNAs expression, but this approach does not include the effects of pre-analytic variables on circulating miRNAs measurement [47,48]. On the other hand, endogenous miRNAs might be considered good reference miRNAs, since their expression is affected by the same variables as the targeted miRNAs. A universally accepted normalization strategy is still lacking. The two main strategies involve the identification of stably expressed reference miRNA previously reported in the literature, or, if miRNA profile has been performed by micro-array or sequencing technologies, the calculation of the global mean expression value of all expressed miRs in a given sample [49]. Thus, the selection of different normalization strategies may affect miRNA quantification and divergence between studies. We used miR-16 as a reference based on previously reported data, being aware that this is one of the many methods that could be used [40,50]. The difficulties associated with hemolysis and platelet contamination of plasma samples are also significant, but it is conceivable that this issue can be mitigated by reducing the degree of red blood cell and platelet-derived miRNA contamination with adequate centrifugation and plasma collection [51]. Other limitations of this study include the small sample size, which should be implemented, and the need for a large validation group.

## Conclusions

The present results lay the basis for a breeding program that will help the CKCS’ breeders in their targeted selection to obtain healthier subjects with a reasonable life expectancy. At the same time, highlighting the risk of the development of the disease at an earlier stage will favour a preventive screening and a mitigating therapeutical approach.

To that end, the identification of early biomarkers for premature MMVD would be a helpful addition.

## Acknowledgements

The authors are grateful to the many dog owners and breeders for their enthusiastic participation to this work and for their availability.

## References

1. Parker HG, Kilroy-Glynn P. Myxomatous mitral valve disease in dogs: does size matter? J Vet Cardiol. 2012;14(1): 19–29.

2. Madsen MB, Olsen LH, Haggstrom J, Höglund K, Ljungvall I, Falk T, et al. Identification of 2 Loci associated with development of myxomatous mitral valve disease in cavalier king charles spaniels. J Hered. 2011;102: S62–S67.

3. Borgarelli M, Haggstrom J. Canine degenerative myxomatous mitral valve disease: natural history, clinical presentation and therapy. Vet Clin North Am Small Anim Pract. 2010;40: 651–663.

4. Serfass P, Chetboul V, Sampedrano CC, Nicolle A, Benalloul T, Laforge H, et al. Retrospective study of 942 small-sized dogs: Prevalence of left apical systolic heart murmur and left-sided heart failure, critical effects of breed and sex. J Vet Cardiol. 2006;8(1): 11–18.

5. Lewis T, Swift S, Woolliams JA, Blott S. Heritability of premature mitral valve disease in Cavalier King Charles spaniels. Vet J. 2011;188(1): 73–76.

6. Swenson L, Haggstrom J, Kvart C. Relationship between parental cardiac status in Cavalier King Charles spaniels and prevalence and severity of chronic valvular disease in offspring. J Am Vet Med Assoc. 1996;208: 2009–2012.

7. Pedersen HD, Lorentzen KA, Kristensen BO. Echocardiographic mitral valve prolapse in cavalier King Charles spaniels: epidemiology and prognostic significance for regurgitation. Vet Rec. 1999;144: 315–320.

8. Gholaminejad A, Zare N, Dana N, Shafie D, Mani A, Javanmard SH. A meta-analysis of microRNA expression profiling studies in heart failure. Heart Fail Rev. 2021;26(4): 997–1021.

9. Duggal B, Gupta MK, Naga Prasad SV. Potential Role of microRNAs in Cardiovascular Disease: Are They up to Their Hype? Curr Cardiol Rev. 2016;12(4): 304–310.

10. Chen X, Ba Y, Ma L, Cai X, Yin Y, Wang K, et al. Characterization of microRNAs in serum: a novel class of biomarkers for diagnosis of cancer and other diseases. Cell Res. 2008;18: 997–1006.

11. Lawrie CH, Gal S, Dunlop HM, Pushkaran B, Liggins AP, Pulford K, et al. Detection of elevated levels of tumour-associated microRNAs in serum of patients with diffuse large B-cell lymphoma. Br J Haematol. 2008;141: 672–675.

12. Mitchell PS, Parkin RK, Kroh EM, Fritz BR, Wyman SK, Pogosova-Agadjanyan EL, et al. Circulating microRNAs as stable blood-based markers for cancer detection. Proc Natl Acad Sci USA. 2008;105: 10513–10518.

13. Matias-Garcia PR, Wilson R, Mussack V, Reischl E, Waldenberger M, Gieger C, et al. Impact of long-term storage and freeze-thawing on eight circulating microRNAs in plasma samples. PLoS One. 2020;15(1): e0227648.

14. Balzano F, Deiana M, Dei Giudici S, Oggiano A, Baralla A, Pasella S, et al. miRNA Stability in Frozen Plasma Samples. Molecules. 2015;20(10): 19030–19040.

15. Paul P, Chakraborty A, Sarkar D, Langthasa M, Rahman M, Bari M, et al. Interplay between miRNAs and human diseases. J Cell Physiol. 2018;233(3): 2007–2018.

16. Dalla Costa E, Dai F, Lecchi C, Ambrogi F, Lebelt D, Stucke D, et al. Towards an improved pain assessment in castrated horses using facial expressions (HGS) and circulating miRNAs. Vet Rec. 2021;188(9): e82.

17. Miretti S, Lecchi C, Ceciliani F, Baratta M. MicroRNAs as Biomarkers for Animal Health and Welfare in Livestock. Front Vet Sci. 2020;18;7: 578193.

18. Lecchi C, Zamarian V, Borriello G, Galiero G, Grilli G, Caniatti M, et al. Identification of Altered miRNAs in Cerumen of Dogs Affected by Otitis Externa. Front Immunol. 2020;29;11: 914.

19. Lecchi C, Zamarian V, Gini C, Avanzini C, Polloni A, Rota Nodari S, et al. Salivary microRNAs are potential biomarkers for the accurate and precise identification of inflammatory response after tail docking and castration in piglets. J Anim Sci. 2020;98(5): skaa153.

20. Corsten MF, Dennert R, Jochems S, Kuznetsova T, Devaux Y, Hofstra L, et al. Circulating MicroRNA208b and MicroRNA-499 reflect myocardial damage in cardiovascular disease. Circ Cardiovasc Genet. 2010;3: 499–506.

21. Fukushima Y, Nakanishi M, Nonogi H, Goto Y, Iwai N. Assessment of plasma miRNAs in congestive heart failure. Circ J. 2011;75: 336–340.

22. Wang GK, Zhu JQ, Zhang JT, Li Q, Li Y, He J, et al. Circulating microRNA: a novel potential biomarker for early diagnosis of acute myocardial infarction in humans. Eur Heart J. 2010;31: 659–666.

23. Ro WB, Kang MH, Song DW, Lee SH, Park HM. Expression Profile of Circulating MicroRNAs in Dogs With Cardiac Hypertrophy: A Pilot Study. Front Vet Sci. 2021; Apr 9;8: 652224.

24. Yang VK, Tai AK, Huh TP, Meola DM, Juhr CM, Robinson NA, et al. Dysregulation of valvular interstitial cell let-7c, miR-17, miR-20a, and miR-30d in naturally occurring canine myxomatous mitral valve disease. PLoS One. 2018;13(1): e0188617.

25. Yang VK, Loughran KA, Meola DM, Juhr CM, Thane KE, Davis AM, et al. Circulating exosome microRNA associated with heart failure secondary to myxomatous mitral valve disease in a naturally occurring canine model. J Extracell Vesicles. 2017;6(1): 1350088.

26. Li Q, Freeman LM, Rush JE, Laflamme DP. Expression Profiling of Circulating MicroRNAs in Canine Myxomatous Mitral Valve Disease. Int J Mol Sci. 2015;16(6): 14098–14108.

27. Lu CC, Liu MM, Culshaw G, Clinton M, Argyle DJ, Corcoran BM. Gene network and canonical pathway analysis in canine myxomatous mitral valve disease: a microarray study. Vet J. 2015;204(1): 23–31.

28. Jung SW, Bohan A. Genome-wide sequencing and quantification of circulating microRNAs for dogs with congestive heart failure secondary to myxomatous mitral valve degeneration. Am J Vet Res. 2018;79(2): 163–169.

29. Keene BW, Atkins CE, Bonagura JD, Fox PR, Häggström J, Fuentes VL, et al. ACVIM consensus guidelines for the diagnosis and treatment of myxomatous mitral valve disease in dogs. J Vet Intern Med. 2019;33(3): 1127–1140.

30. Rishniw M. Murmur grading in humans and animals: past and present. J Vet Cardiol. 2018;20(4): 223–233.

31. Brown S, Atkins C, Bagley R, Carr A, Cowgill L, Davidson M, et al. American College of Veterinary Internal Medicine. Guidelines for the identification, evaluation, and management of systemic hypertension in dogs and cats. J Vet Intern Med. 2007;21(3): 542–558.

32. Acierno MJ, Brown S, Coleman AE, Jepson RE, Papich M, Stepien RL, et al. ACVIM consensus statement: Guidelines for the identification, evaluation, and management of systemic hypertension in dogs and cats. J Vet Intern Med. 2018;32(6): 1803–1822.

33. Thomas WP, Gaber CE, Jacobs GJ, Kaplan PM, Lombard CW, Moise NS, et al. Recommendations for standards in transthoracic two-dimensional echocardiography in the dog and cat. J Vet Intern Med. 1993;7: 247–252.

34. Chetboul V, Tissier R. Echocardiographic assessment of canine degenerative mitral valve disease. J Vet Card. 2012;14(1): 127–148.

35. Ponikowski P, Voors AA, Anker SD, Bueno H, Cleland JG, Coats AJ, et al. Authors/Task Force Members; Document Reviewers. 2016 ESC Guidelines for the diagnosis and treatment of acute and chronic heart failure: The Task Force for the diagnosis and treatment of acute and chronic heart failure of the European Society of Cardiology (ESC). Eur Heart J. 2016;37(27): 2129–2200.

36. Grimes JA, Prasad N, Levy S, Cattley R, Lindley S, Boothe HW, et al. A comparison of microRNA expression profiles from splenic hemangiosarcoma, splenic nodular hyperplasia, and normal spleens of dogs. BMC Vet Res. 2016;12: 272–284.

37. Zhao FR, Su S, Zhou DH, Zhou P, Xu TC, Zhang LQ, et al. Comparative analysis of microRNAs from the lungs and trachea of dogs (Canis familiaris) infected with canine influenza virus. Infect Genet Evol. 2014;21: 367–374.

38. Bustin SA, Benes V, Garson JA, Hellemans J, Huggett J, Kubista M, et al. The MIQE guidelines: Minimum information for publication of quantitative real-time PCR experiments. Clin Chem. 2009;55(4): 611–622.

39. Mussack V, Hermann S, Buschmann D, Kirchner B, Pfaffl MW. MIQE-Compliant Validation of MicroRNA Biomarker Signatures Established by Small RNA Sequencing. In: Biassoni R, Raso A, editors. Quantitative Real-Time PCR. Methods in Molecular Biology, vol 2065. Humana, New York, NY.

40. Hulanicka M, Garncarz M, Parzeniecka-Jaworska M, Jank M. Plasma miRNAs as potential biomarkers of chronic degenerative valvular disease in Dachshunds. BMC Vet Res. 2014;10: 205.

41. Šimundić AM. Measures of Diagnostic Accuracy: Basic Definitions. EJIFCC. 2009;19(4): 203–211.

42. Borgarelli M, Buchanan JW. Historical review, epidemiology and natural history of degenerative mitral valve disease. J Vet Cardiol. 2012;14(1): 93–101.

43. Zhang X, Dong S, Jia Q, Zhang A, Li Y, Zhu Y, et al. The microRNA in ventricular remodeling: the miR-30 family. Biosci Rep. 2019;39(8): BSR20190788.

44. Kim JO, Park JH, Kim T, Hong SE, Lee JY, Nho KJ, et al. A novel system-level approach using RNA-sequencing data identifies miR-30-5p and miR-142a-5p as key regulators of apoptosis in myocardial infarction. Sci Rep. 2018;8(1): 14638.

45. Miranda K, Yang X, Bam M, Murphy EA, Nagarkatti PS, Nagarkatti M. MicroRNA-30 modulates metabolic inflammation by regulating Notch signaling in adipose tissue macrophages. Int J Obes. 2018;42(6): 1140–1150.

46. Efthimiadis G, Tzimagiorgis G. A critical approach for successful use of circulating microRNAs as biomarkers in cardiovascular diseases: the case of hypertrophic cardiomyopathy. Heart Fail Rev 2022; 27(1): 281–294.

47. McDonald JS, Milosevic D, Reddi HV, Grebe SK, Algeciras-Schimnich A. Analysis of circulating microRNA: preanalytical and analytical challenges. Clin Chem. 2011;57: 833–840.

48. Cheng HH, Yi HS, Kim Y, Kroh EM, Chien JW, Eaton KD, et al. Plasma processing conditions substantially influence circulating microRNA biomarker levels. PLoS One 2013;8(6): e64795.

49. Navickas R, Gal D, Laucevičius A, Taparauskaitė A, Zdanytė M, Holvoet P. Identifying circulating microRNAs as biomarkers of cardiovascular disease: a systematic review. Card Res. 2016;111(4): 322–337.

50. Noszczyk-Nowak A, Zacharski M Michałek. Screening for circulating miR-208a and - b in different cardiac arrhythmias of dogs. J Vet Res. 2018;62(3): 359–363.

51. Kirschner MB, Edelman JJ, Kao SC, Vallely MP, van Zandwijk N, Reid G. The impact of hemolysis on cell-free microRNA biomarkers. Front Genet. 2013;4: 94.

